# Investigating the concept of accessibility for predicting novel RNA-RNA interactions

**DOI:** 10.1101/2021.06.03.446902

**Authors:** Sabine Reißer, Irmtraud M Meyer

## Abstract

State-of-the-art methods for predicting novel *trans* RNA-RNA interactions use the so-called accessibility as key concept. It estimates whether a region in a given RNA sequence is accessible for forming *trans* interactions, using a thermodynamic model which quantifies its secondary structure features. RNA-RNA interactions are then predicted by finding the minimum free energy base-pairing between the two transcripts, taking into account the accessibility as energy penalty.

We investigated the underlying assumptions of this approach using the two methods RNAplex and IntaRNA on two datasets, containing sRNA-mRNA and snoRNA-rRNA interactions, respectively.

We find that (1) known *trans* RNA-RNA interactions frequently overlap regions containing RNA structure features, (2) the estimated accessibility reflects sRNA structures fairly well, but often disagrees with structure annotations of longer transcripts, (3) the prediction performance of RNA-RNA interaction prediction methods is independent of the quality of the estimated accessibility profiles, and (4) one important overall effect of accessibility profiles is to prevent the thermodynamic model from predicting too long interactions.

Based on our findings, we conclude that the accessibility concept to the minimum free energy approach to predicting novel RNA-RNA interactions has conceptual limitations and discuss potential ways of improving the field in the future.

## INTRODUCTION

Direct *trans* RNA-RNA interactions between two transcripts are key to mediating many biological mechanisms in diverse living organisms (1, 2, 3, 4, 5). SnRNAs bind to nascent RNA transcripts to guide splicing into mature transcripts (1). SnoRNAs play an important role during ribosome biogenesis, enabling chemical modifications like methylation and pseudouridylation of bases which are key to the ribosome’s correct functioning (6, 7). Also the codon-anticodon recognition of tRNAs is facilitated by RNA-RNA interactions (8). In eukaryotes, miRNAs regulate gene expression *via* RNA interference, by binding to mRNAs and thereby blocking translation (2). In a similar way, sRNAs can block translation of bacterial mRNAs (3, 4).

In the following, *trans* RNA-RNA interactions and *trans* base pairs will refer to base pairs formed between two transcripts, i.e. inter-molecular base pairs, whereas *cis* RNA-RNA interactions and cis base pairs will denote intramolecular base pairs within the same transcript, i.e. features of the transcript’s RNA secondary structure.

There is a plethora of published tools for the target prediction of specific query molecules, like miRNA (9, 10, 11, 12, 13, 14, 15, 16, 17) and sRNA (18, 19, 20, 21, 22), but also for target prediction of C/D box snoRNA (23), H/ACA box snoRNAs (24), siRNAs (25) and piRNAs (26). In order to be able to discover novel classes of *trans* RNA-RNA interactions, however, we require computational methods that are able to discover entirely new biological types of interactions whose details, i.e. *cis* and *trans* base pairs, are not yet known.

For predicting these novel biological classes of *trans* RNA-RNA interactions, there exist dedicated computational methods, for the most recent reviews see (27, 28). These so-called *ab initio* methods can be subdivided into three broad classes. First, non-comparative methods that take only the two transcripts of interest as input, e.g. IntaRNA (29, 30), RNAplex (31), RNAduplex (32), RIsearch (33), RNAup (34) and RNAcofold (35). Second, comparative methods that utilise two multiple sequence alignments (MSAs) as input (one MSA for each transcript of interest), e.g. RNAplex (31), PetCofold (36) and RNAaliduplex (32). And, third, alignment-free, comparative methods which take two sets of unaligned, orthologous sequences as input (one set for each transcript of interest) e.g. IRBIS (37).

Many transcripts that are known to interact *in vivo* with another transcript in *trans* may also exhibit RNA structure, i.e. *cis* RNA-RNA interactions, at the same or a different time of their cellular life. In H/ACA snoRNA-rRNA interactions, for example, a distinct snoRNA secondary structure is required for the *trans* RNA-RNA interaction to form so the rRNA can be correctly pseudouridylated. The rRNA, however, is also known to exhibit a very distinct RNA secondary structure at a different stage of its cellular life, namely while being part of the mature ribosome (38, 39). Another example illustrating the complexities of *cis* and *trans* interactions during a transcript’s life in *vivo* is the *trans* interaction between the sRNA OxyS and the mRNA *fhlA* in *E. coli* where both transcripts on their own exhibit distinct secondary structure features in direct proximity to the known *trans* interaction site (3).

Right now, the state-of-the-art in terms of prediction accuracy for novel *trans* RNA-RNA interactions is obtained by computational methods that work in a non-comparative way. These consider as input only the sequences of the two transcripts of interest and assume that *trans* RNA-RNA interactions are more likely to form in regions of the transcript that are devoid of RNA structure features. To this end, they estimate the so-called accessibility along each input sequence using a minimum free energy (MFE) strategy. For this, the entire input sequence (i.e. the transcript of interest) is considered to be in thermodynamic equilibrium in solution without any *trans* interaction partners and all potential, pseudo-knot-free RNA secondary structures are approximately estimated to quantify which regions along the sequence are more devoid of RNA structure features than others.

RNAup (34), AccessFold (40), and RIblast (41) calculate the partition function of secondary structure features for the entire input sequence, while RIblast uses an additional parameter to limit the maximum base pair span. To keep the calculation computationally tractable, RNAplfold uses both a constraint on the maximum base pair span and a window of length *W* (shorter or equal the sequence length), which is moved along the sequence, and local RNA structure features are calculated for the sub-sequence inside the window. Based on the free energy of these RNA structure features, base pairs in these windows are assigned a probability to be formed in thermodynamic equilibrium. The global probability for a specific base pair is then obtained by averaging over all windows containing the base pair. From this, the probability for each sequence position (or stretch of positions) to be unpaired is derived, which is its final ‘accessibility’ (42, 43).

Once the accessibilities along both input sequences have been estimated, the *trans* RNA-RNA interactions are typically predicted using a thermodynamic approach which essentially captures the assumption that the two transcripts are in thermodynamic equilibrium and that they aim to settle in the joint configuration with the smallest overall Gibbs free energy, penalised by the ‘opening energies’ derived from the probability of the two binding regions to be unpaired. In other words, these methods aim to predict the MFE *trans* RNA-RNA configuration between the two input RNA sequences (29, 31).

Even the best state-of-the-art methods, however, have trouble generating high-quality predictions when the two RNA sequences are anything but rather short (27, 28).

On the experimental side, a number of novel high-throughput methods have been recently published, which are able to capture both *cis* and *trans* RNA-RNA interactions *in vivo* and on a transcriptome-wide scale, namely PARIS, SPLASH, and LIGR-seq (44, 45, 46). The raw data generated by these methods comes in terms of so-called duplexes, where one duplex corresponds to a single pair of either *cis* or *trans* interacting sub-sequences that have been cross-linked into the same chimeric read as part of the experimental procedure. These exciting new methods, however, are still in their infancy as (1) the probing compound psoralen (or the psoralen derivative AMT in case of LIGR-seq) has biases since it only covalently cross-links stacked pyrimidines on opposite strands (implying that perfectly stable helices composed only of {*G*,*C*} base pairs will not be crosslinked and thus not detected) and (2) the overall probing of the *cis* and *trans* interactome is generally not deep enough due to several efficiency bottle-necks (47).

This implies that computational methods for detecting novel *trans* RNA-RNA interactions based on transcript sequence information alone are still very much needed, not only to explore the universe of potential *cis* and *trans* interactions within many readily available transcriptome datasets, but particularly to generate hypotheses on potential interactions partners and corresponding *cis* and *trans* features that can then be experimentally validated in dedicated experiments.

In order to improve the current state-of-the-art in the field of predicting novel *trans* RNA-RNA interactions, we were thus keen to investigate (1) whether the underlying assumption, namely that regions of potential *trans* RNA-RNA interactions have to be devoid of RNA structure features, is justified and (2) whether the commonly used computational method RNAplfold is capable of estimating accessibility correctly. We further analyse how different settings in the accessibility calculation influence both the accuracy of the accessibility profiles and the RNA-RNA interaction prediction performance for the two state-of-the-art programs, IntaRNA and RNAplex. Finally, we investigate the differences in prediction with and without the use of accessibility profiles.

Our results show that (1) contrary to the commonly made assumption, known *trans* RNA-RNA interactions frequently overlap regions that are known to also contain RNA structure features, (2) the estimated accessibility reflects sRNA structure annotations fairly well, but often disagrees with structure annotations of longer transcripts, (3) the prediction performance of RNA-RNA interactions prediction software is independent of the quality of the estimated accessibility profiles, and (4) one important overall effect of considering accessibility profiles is to prevent the thermodynamic model from predicting too long interactions.

The manuscript is structured in the following way: in the section ‘DATASETS’, we describe the two datasets investigated in our study which represent two distinct biological classes, sRNA-mRNA and snoRNA-rRNA interactions. In ‘METHODS’, we describe the full details of our computational analysis. The ‘RESULTS’ section is then structured to support each of the four findings listed above. Finally, we discuss our findings and conclude.

## DATASETS

### sRNA - mRNA

This dataset comprises 109 experimentally verified sRNA-mRNA interactions which have been previously published in the survey (27). Of those, 64 interactions are from Escherichia coli str. K-12 substr. MG1655 (*E. coli*), and 45 from Salmonella enterica subsp. enterica serovar Typhimurium str. LT2 (*S. enterica*). The average RNA-RNA interaction duplex length is 21 nt which contain on average of 7.3 unpaired nucleotides, i.e. nucleotides within bulges or loops. The characteristic features of this dataset such as the distribution of interaction lengths and the number of mismatches/bulges per RNA-RNA interaction are shown in Supplementary Material Figure S1A-C. All mRNAs contain the 5’ untranslated regions (5’UTRs) extending to the stop codon of the next gene upstream, or 300 nt if the next gene is at a larger distance. Compared to (27), we replaced three genes, because the RNA-RNA interaction is located in the coding sequence of the next gene upstream, and not in the 5’UTR of the originally specified gene. Thus, rpoS, ilvE, and yigL were replaced by nlpD, ilvM, and pldB, respectively. We also updated the genomic coordinates of *S. enterica* to correctly reflect the reference genome NC_003197.2.

It is worthy to note that there are 5 sRNA-mRNA pairs for which two interaction sites are known, namely the pairs with ids 26/27, 43/44, 56/57, 71/72, 73/74. The complete dataset can be found in the Supplementary Material File sRNA-mRNA_RRI_RSS.csv. Note that column ‘srna_sec_str_source’ (origin of sRNA structure annotation) contains either ‘RFAM’ or the PMID of the corresponding publication.

#### Structural annotations of sRNAs

The structural annotation of this dataset contains the known RNA secondary structures of all sRNAs which either derive from published data (generated from experimental data, or predicted with comparative or MFE approaches), or have been predicted *ab initio* by us. Structures generated *via* the comparative approach have been taken from the RFAM database (48). For two sRNAs, RNA secondary structures were predicted by us using the program RNAfold from ViennaRNA v.2.4.16, with default settings (32). This can be justified as this MFE-based method can be expected to perform well for these rather short sequences (*S. enterica* SgrS, 239 nt, *S. enterica* ChiX, 81 nt).

#### Structural annotations of mRNAs

For 11 (out of 90) mRNAs, published RNA secondary structures derived from SHAPE-MaP data covering the full transcript were available in the Supplemental Material of (49). To obtain the structures contained in the region of interest, we excised the relevant sub-structure using *RNAtools.py* which is part of Superfold (https://github.com/Weeks-UNC/Superfold) (50). We considered the conserved motifs shown in (50) as well as the RNA secondary structures from the cell-free and the in-cell environment for our analysis, all structure annotations can be found in the Supplementary Material File sRNA-mRNA_RRI_RSS.csv, listed under ‘alternative’. For 5 mRNAs, other experimental data was available, as noted in the same file.

Most mRNAs (75 out of 90) in the dataset were missing any structural annotation and had also no published SHAPE-MaP data. We annotated those using the same procedure as in (49) for the prediction of mRNA structures using SHAPE-MaP data as restraints, but using unknown SHAPE values (set to “no-data” (nan)). This involved the use of Superfold v.1.1 together with RNAstructure v.6.2 (50, 51). Parameters for SuperFold were: SHAPEslope = 1.8, SHAPEintercept = −0.6, trimInterior = 300, partitionWindowSize = 1500, partitionStepSize = 100, foldWindowSize = 3000, foldStepSize = 300, maxPairingDist = 500.

To summarise, as only few mRNA structures come with experimental evidence and as most were predicted *in silico*, we place only limited trust in the structural annotation of the mRNAs and thus show the relevant results only in the Supplementary Material.

### snoRNA - rRNA

This dataset comprises 52 verified RNA-RNA interactions between pairs of snoRNAs and ribosomal RNA (rRNA) from *Saccharomyces cerevisiae S288c* (*S. cerevisiae*), 18 of which are with the small subunit rRNA 18S and 34 with the large subunit rRNA 25S. The average length of an RNA-RNA interaction duplex is 13 nt, with on average 0.54 nucleotides in bulges or loops. Please refer to Supplementary Material Figure S1D-F for the detailed characteristics of the dataset such as the distribution of interaction lengths and the number of mismatches/bulges per RNA-RNA interaction.

It is interesting to note that also in this dataset, there are 5 snoRNA-rRNA pairs for which there are two interaction sites, namely the pairs with ids 1/2, 5/6, 22/23, 35/36, 43/44. The complete dataset can be found in Supplementary Material File snoRNA-rRNA_RRI_RSS.csv.

#### Structural annotations of rRNAs

The secondary structures of *S. cerevisiae* 18S and 25S ribosomal subunits have been obtained from (39).

#### Structural annotations of snoRNAs

The annotation of this dataset comprises RNA secondary structures for all snoRNAs. The RNA secondary structures have been taken from the RFAM database where present (48). RNA secondary structures missing for 5 snoRNAs in RFAM were predicted by us using the program RNAfold from ViennaRNA v.2.4.16 with default settings (32). It is reasonable to expect this MFE-based method to perform well for these rather short sequences (78-98 nt).

Overall, there is almost no experimental evidence for the annotated RNA secondary structures of the snoRNAs. The RNA secondary structures of RFAM are based on evolutionary evidence in terms of covariation from an underlying multiple sequence alignment (MSA). These MSAs, however, are fairly heterogeneous in terms of the number and the pairwise similarity of the underlying sequences. It is worthy to note that many of the official RFAM structures have almost no base pairs at all. We therefore place only limited trust in these snoRNA structures and show the corresponding figures in the Supplementary Material.

## METHODS

In the following, ‘query’ refers to the shorter of the two interacting transcripts, i.e. either sRNA or snoRNA and ‘target’ to the longer transcript of the pairwise interaction, i.e. either an mRNA or rRNA, depending on the dataset.

### Analysis of conflicts between RNA secondary structure and RNA-RNA interaction

One of our key goals is to investigate the validity of the accessibility strategy of the state-of-the-art methods. This strategy assumes that regions involved in *trans* interactions should be devoid of RNA secondary structures features. We thus define a *conflict* as a situation, where a nucleotide that is known to form a *trans* base pair is also known to form a *cis* base pair as part of the transcript’s RNA secondary structure.

We analysed conflicts between RNA secondary structure and *trans* RNA-RNA interaction base pairs separately for query and target, using

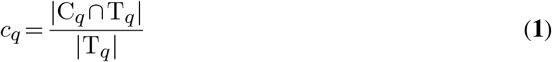

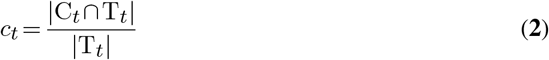

Here, C_*q*_ denotes the set of query nucleotides that are known to be base-paired in *cis* (i.e. the nucleotides which are base-paired in the transcript’s known RNA secondary structure) and T_*q*_ the set of query nucleotides that are known to be base-paired in *trans*. This implies that *c_q_* is the fraction of *trans* nucleotides that are also paired in *cis* in the query. In a similar way, *c_t_* denotes the fraction of *trans* nucleotides that are also paired in *cis* in the target.

In case of multiple known structure annotations for one transcript (e.g. based on different experimental conditions), we use *c_q/t_* to denote the average value. For the query-target pairs with two known RNA-RNA interactions each, *c_q/t_* is calculated for the combination of both RNA-RNA interactions.

### RNA-RNA interaction prediction tools

The two state-of-the-art methods for predicting *trans* interactions that we consider, IntaRNA and RNAplex, both use RNAplfold from the ViennaRNA package (32) for estimating the accessibilities along any given RNA sequence.

We employ RNAplfold from ViennaRNA v.2.4.15 with the following settings to estimate the accessibilities for query and target sequences: -W *W* -L *L* -u *u* -O. Here, *L* denotes the maximum distance of two base-paired nucleotides inside the sliding window of size *W*, i.e. *L* ≤ *W*. This window is moved along the sequence to calculate averaged base pairing probabilities. The values for *W* and *L* are specified in the text. *u* denotes the maximum length (in nucleotides) of any region for which unpaired probabilities are calculated and reported in the output. For *u* = 10, for example, the output of RNAplfold contains the probabilities for all possible subsequences between 1 and 10 nucleotides length to be unpaired. We used as default value *u* = 60, unless stated otherwise. “-O” is specified to convert unpaired probabilities to energy penalties in the output, the output files then have the suffix ‘_openen’.

We use IntaRNA v.3.2.0, with energy penalties pre-calculated by RNAplfold as specified above, using the following settings (29, 30): --qAcc E --qAccFile qname_openen --tAcc E --tAccFile tname_openen -q qname.fa -t tname.fa --outMode C. This is equivalent to running IntaRNA with --qAccW *W* --qAccL *L* --tAccW *W* --tAccL *L* --intLenMax *u*, as IntaRNA invokes RNAplfold to calculate the required accessibilities. This means that for IntaRNA, the maximum interaction length is 60 nucleotides, which theoretically covers all possible known interactions in our two datasets and is the recommended setting for sRNA target prediction (52). Note that specific recommendations for improved sRNA target prediction performance have been published recently (52).

As second state-of-the-art program for predicting *trans* interactions, we use RNAplex from ViennaRNA v.2.4.15 with the following parameters (32): -q qname.fa -t tname.fa -a acc -f 2. Here, ‘acc’ specifies the name of the folder containing qname_openen and tname_openen. We use the tag ‘-f 2’ (fast approximate energy model with re-computation of the actual interaction energies) on recommendation of the authors of RNAplex since the default option for the backtracking (‘-f 0’) results in the prediction of non-canonical base pairs. For RNAplex, no maximum interaction length can be set, since the option “-l” (maximal length of an interaction) is ignored by the program.

### Measuring the quality of the accessibility estimation

Accessibilities for query and target are calculated using RNAplfold as above but without “-O”, yielding unpaired probabilities per nucleotide as output, instead of opening energies. Single nucleotide unpaired probabilities (column ‘l=1’) are used to analyse accessibility. In order to quantify, how much the estimated accessibility agrees with the known RNA secondary structure annotation, we define the *quality of an accessibility profile Q* as follows:

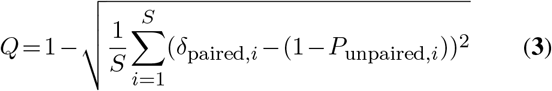

where *S* denotes the sequence length in nucleotides, *i* the sequence position, *δ*_paired,*i*_ (1, if paired, 0, if unpaired) the pairing status of sequence position *i* according to the known RNA secondary structure annotation, and *P*_unpaired, *i*_ the estimated probability of sequence position *i* being unpaired. In a nutshell, *Q* ∈ [0, 1] with *Q* = 1 if the accessibility profile perfectly reflects the known structural annotation and *Q* = 0 if both are in complete disagreement along the entire transcript.

### Prediction performance

For both IntaRNA and RNAplex, we take the MFE prediction for each query-target pair to judge the prediction performance. As is common, we measure the prediction performance in terms of sensitivity (defined as Sens = TP/(TP+FN)), the positive predicted value (precision, PPV = TP/(TP + FP)), and the F1 score (F1=2· (Sens· PPV)/(Sens + PPV)), which is the harmonic mean of sensitivity and PPV. As usual, *TP* (true positives) denotes the number of correctly predicted base pairs, *FP* (false positives) the number of incorrectly predicted base pairs and *FN* (false negatives) the number of base pairs present in the reference, but missing in the prediction. A high sensitivity thus implies that most known base pairs were correctly predicted, whereas a high *PPV* means that there are few predicted base pairs that do not coincide with known ones.

## RESULTS

### Known *trans* RNA-RNA interactions frequently overlap regions with known RNA structure features

The implicit assumption underlying the accessibility-based approach to *trans* RNA-RNA interaction prediction is that potential binding sites are essentially devoid of RNA structure features or – in other words – that there are no conflicting *cis* and *trans* base pairs. To test the validity of this assumption, we first systematically calculated *cis*/*trans* conflicts in both of our datasets and we find that there is a significant amount of conflicts in both datasets.

Figure 1 shows the degree of conflicts between known sRNA secondary structures and known sRNA-mRNA interactions, *c*_sRNA_. While there are many green squares, indicating that the *trans*-interacting region is essentially unstructured, there are also a considerable number of known *trans* RNA-RNA interaction sites which are also known to be base-paired in *cis* at some stage of the transcript’s cellular life. Spf, FnrS, and DsrA sRNAs have the largest overlaps of 67-75%. For mRNAs, there are many more conflicts between known *cis* and known *trans* base pairs, as can be seen from the large amount of yellow-magenta fields in Supplementary Material Figure S2. One has to keep in mind that most of the mRNA structures have been predicted *in silico* using an MFE method and we have limited trust in them. Nevertheless, these transcripts are so long that it can be expected that they exhibit complex RNA secondary structure features, making conflicts between *cis* and *trans* base pairs likely, regardless of the exact structure. The amount of conflicts in sRNAs is not correlated with the prediction performance (Pearson correlation *r* = 0.02 between conflict score and F1 score for both IntaRNA and RNAplex, run with setting ‘RNAplex’). The amount of conflicts in mRNAs is slightly negatively correlated with the performance (r = −0.16 for RNAplex, and *r* = −0.25 for IntaRNA), showing that the prediction performance is decreasing slightly whith the amount of conflicting base pairs.

**Figure 1.**
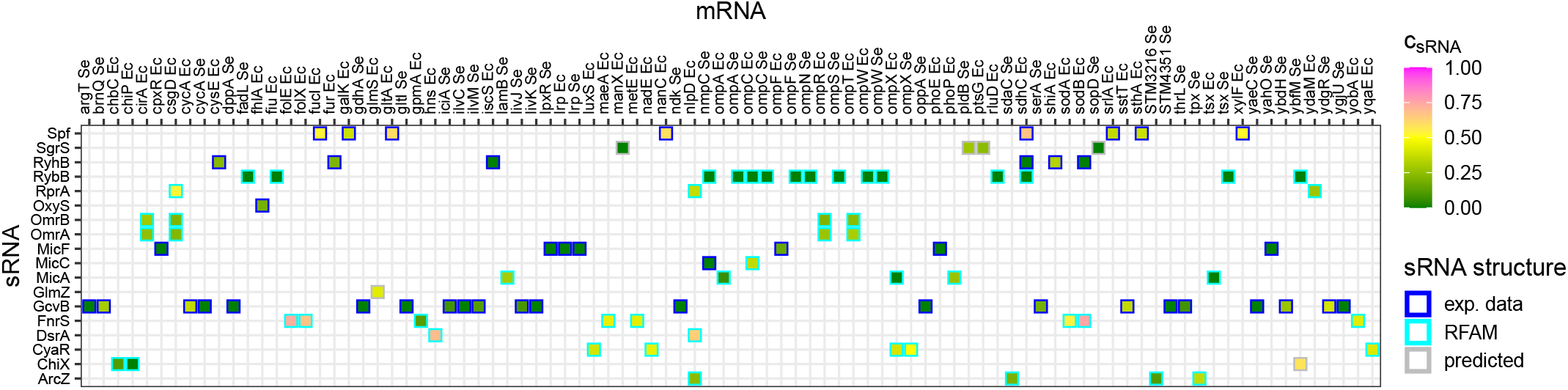
Overview of the amount of conflicts between known sRNA secondary structure and known *trans* RNA-RNA interactions with mRNAs. A value of *c* = 0 implies that there are no conflicting base pairs, whereas a value of *c* = 1 means that all known *trans*-base pairs are in conflict with known sRNA *cis*-base pairs. In the mRNA names, ‘Ec’ refers to *E. coli*, and ‘Se’ refers to *S. enterica*.

Figure 2 shows the RNA secondary structure of the sRNA RprA and the mRNA *csgD* and their interaction as R-chie plot (53). Arcs, i.e. semi-circles, and straight lines in magenta colour indicate base pairs with *cis*/*trans* conflicts.

**Figure 2.**
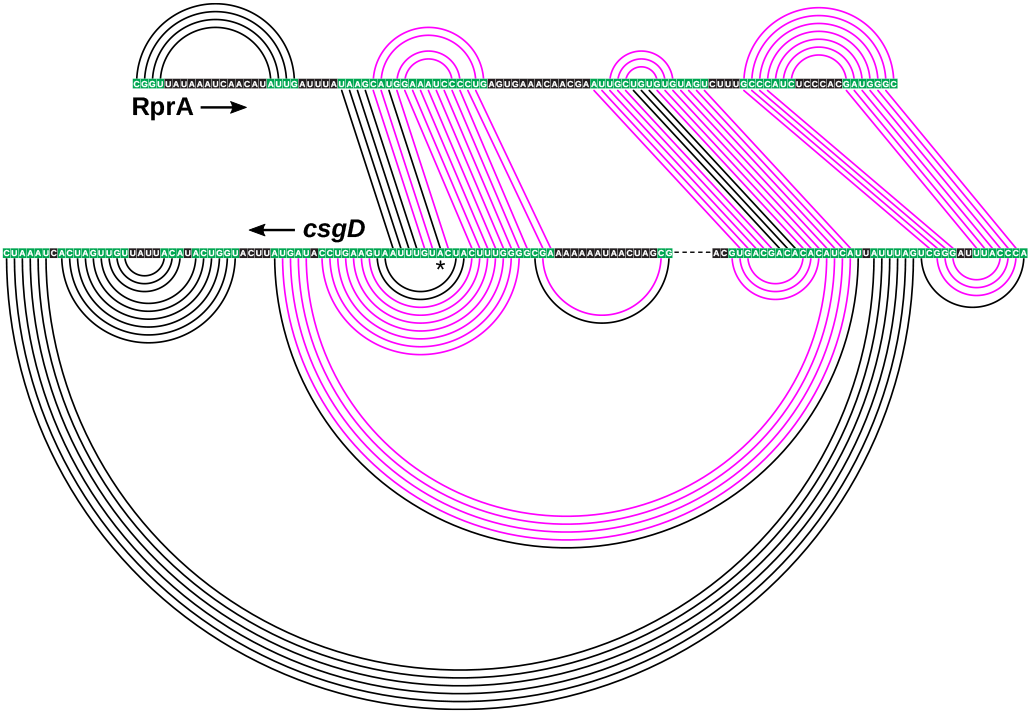
RNA secondary structure and RNA-RNA interactions for the example of sRNA-mRNA pair RprA-*csgD*. Known base pairs with *cis-trans* conflicts are shown in magenta. RNA structures have been trimmed at both ends for better visibility. A green base color indicates standard Watson-Crick base pairing. A dashed line indicates that part of the sequence has been omitted from the figure for visualisation purposes. The arrows indicate 5’ to 3’ direction. The star indicates the position of the start codon.

Figure 3 shows the degree of conflicts between rRNA secondary structure and snoRNA-rRNA interactions, *c*_rRNA_. Both the 18S and 25S ribosomal subunit have well established, complex RNA structures with many helices. The figure shows a high degree of *cis/trans* conflicts in almost all known *trans* RNA-RNA interactions. In many cases, more than 50% of the *trans*-pairing bases are also known to be *cis* base-paired at some stage of the transcript’s cellular life. Supplementary Material Figure S3 shows the conflicts from the perspective of the snoRNAs, *c*_snoRNA_. There are only a few conflicts here, owing to the very low amount of base pairs in the snoRNA structures presented on the RFAM database.

**Figure 3.**
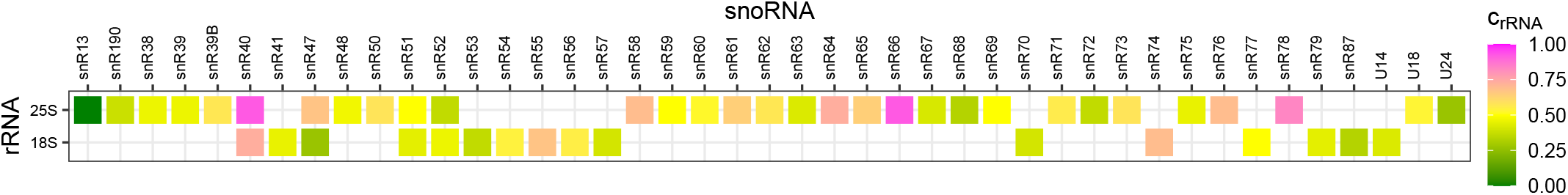
Overview of the amount of conflicts between known rRNA secondary structure and known *trans* RNA-RNA interactions with snoRNAs. As before, *c* = 0 implies that there are no conflicting base pairs, whereas *c* = 1 means that all known *trans*-base pairs are in conflict with known rRNA *cis*-base pairs.

Figure 4 shows the RNA secondary structure and RNA-RNA interaction of three snoRNAs binding to the 25S ribosomal subunit. Again, magenta-coloured arcs and straight lines demonstrate the high amount of conflict between *cis* and *trans* base pairs.

**Figure 4.**
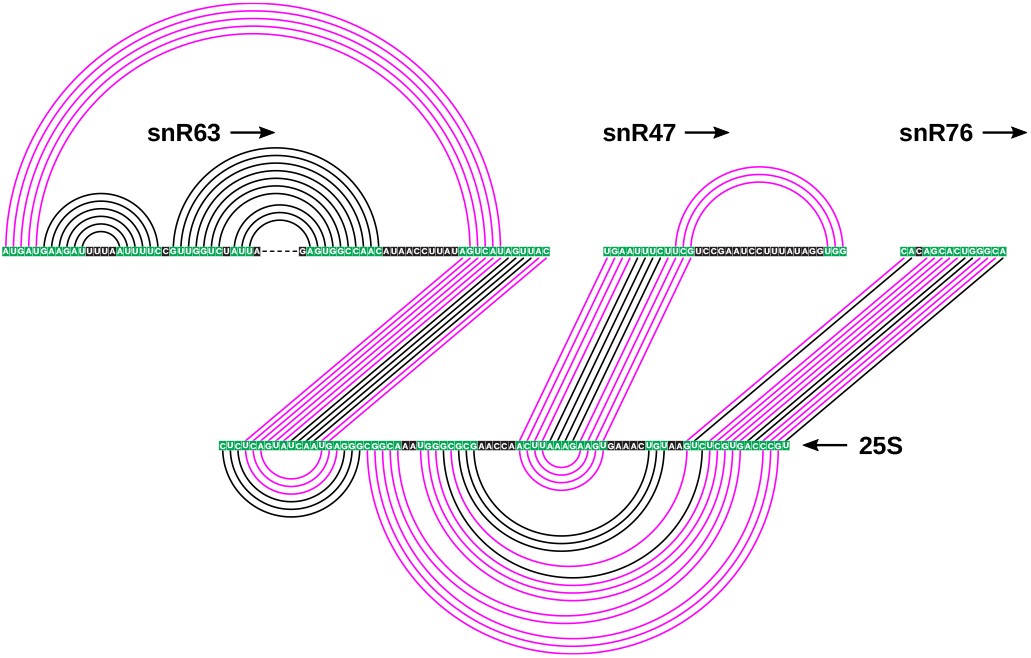
RNA secondary structure and RNA-RNA interactions for the example of snoRNAs snR63, snR47 and snR76, interacting with helices 68 and 69 of the 25S ribosomal subunit it *S. cerevisiae*. Known base pairs with *cis-trans* conflicts are shown in magenta. Once again, we trimmed the RNA structures at both ends for better visibility. A green base color indicates standard Watson-Crick base pairing, whereas a dashed line indicates that part of the sequence has been omitted from the figure for visualisation purposes. The arrows indicate 5’ to 3’ direction.

### The estimated accessibility reflects sRNA structures fairly well, but often disagrees with structures of longer transcripts

RNAplfold is used by both RNA-RNA interaction prediction programs RNAplex and IntaRNA to estimate the accessibility profiles along input transcripts, employing a thermodynamic strategy. A window of *W* nucleotides width is moved along the sequence and local RNA structure features with a maximum base pair span of *L* (*L* ≤ *W*) nucleotides are predicted for the sub-sequence inside the window. This window-based approach is primarily employed to reduce the time and memory requirements of the accessibility estimation, especially for long input sequences. Based on the free energies of the RNA structure features, base pairs in these windows are assigned a probability according to their relative frequency in the corresponding Boltzmann distribution (of pseudoknot free RNA secondary structures for that sequence in thermodynamic equilibrium). The overall probability for a specific base pair is then obtained by averaging over all windows with secondary structure features containing that base pair. From this, the accessibility is derived, which is the probability that a specific sequence position or subsequence is unpaired.

We define the quality measure *Q*, as explained in ‘Methods’, which quantifies the concordance between the estimated unpaired probabilities and the base-paired positions of the known reference RNA secondary structure in the sequence. *Q* = 1 if both are in perfect agreement, and *Q* = 0 if there is complete disagreement. Figure 5 shows two examples of the accessibility profile compared to the known RNA secondary structure, (A), the rather short transcript of sRNA DsrA, where the agreement between calculated accessibility and reference secondary structure is very good (*Q* = 0.85), and (B), the comparatively long transcript of the 18S ribosomal subunit, where the agreement is mediocre (*Q* = 0.55).

**Figure 5.**
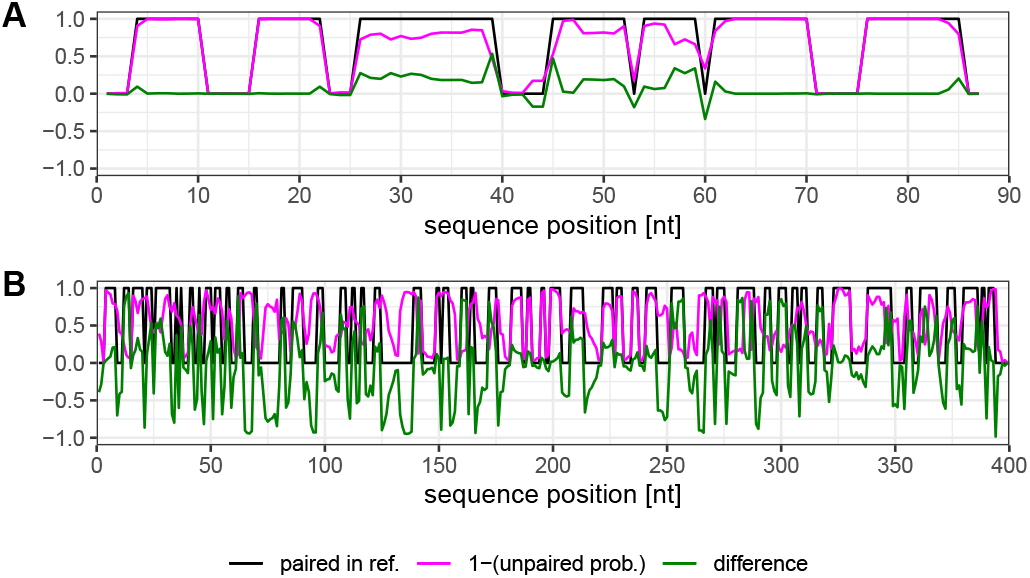
Accessibility profile (in terms of unpaired probabilities for each nucleotide in the transcript) calculated by RNAplfold (magenta) versus known ‘true’ base-paired positions (1: base-paired, 0: unpaired, black). The difference between (1 – unpaired probability) and true base-paired positions is shown in green. (A) sRNA DsrA (RNAplfold settings *W* = *L* = 87), the overall accessibility quality *Q* = 0.85. (B) nucleotides 1-400 of 18S ribosomal subunit of *S. cerevisiae* (RNAplfold settings *W* = *L* = 120). For the shown subsequence of 400 nt, *Q* = 0.54 (for the whole 1800 nt transcript, *Q* = 0.55).

We systematically calculated *Q* for different values of *W* (with *L* = *W*), for all four classes of molecules. Figure 6 shows *Q* as function of *W/S*, where *S* is the length of the sequence, for (A) all sRNAs, and (B) the two ribosomal RNAs. Figure 6A shows that for the short sRNAs, the accessibility calculated with *W* = *S* generally gives the highest *Q* values, i.e. agrees best with the reference structure. The green outlier which starts with a high value of *Q* and decreases with increasing *W* refers to SgrS, for which the reference structure is limited to two hairpins located at the 3’ end of the transcript, due to details of the experimental probing. The rest of the transcript may also be more structured in reality which is likely the reason that *Q* drops with increasing *W*. The average *Q* for *W/S* = 1 is 0.64 (0.65 without SgrS). The mRNA accessibility quality with respect to the reference RNA secondary structure is shown in Supplementary Material Figure S4B. Interestingly, for mRNAs, *Q* reaches also good values and generally increases continuously with increasing *W*, reaching an average *Q* for *W/S* =1 of 0.61 (0.63 without the outliers which are due to incomplete knowledge of the RNA structure where only a part of the sequence was probed, and a large part of the reference structure is without any base pairs). One has to keep in mind, however, that all complete mRNA structures have been computationally derived using a thermodynamic MFE model, and only for a few of them experimental SHAPE data were used as experimental evidence. This means that both the accessibility profiles and the structural annotation have been predicted using the same underlying approach, namely an MFE method. Supplementary Material Figure S4B thus gives us an idea of how well the accessibility, which is an ensemble quality since it reflects all possible RNA structure features, can agree with a single MFE structure of a long transcript such as mRNA.

**Figure 6.**
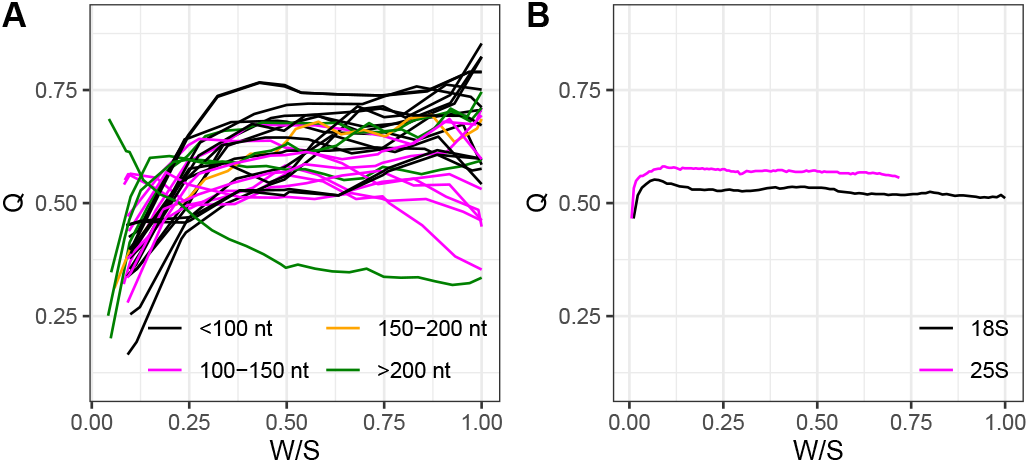
Quality *Q* of the accessibility profiles, as function of the RNAplfold parameters *W* and *L*, with *W* = *L*. *S* denotes the total sequence length. (A) All 27 sRNAs from the sRNA-mRNA dataset, grouped by length. (B) 18S and 25S ribosomal subunits, snoRNA-rRNA dataset.

For rRNAs, whose structures are experimentally well-defined, the highest value of *Q* is reached at *W* = *L* = 120 with 0.55 for 18S and at *W* = *L* = 300 with 0.58 for the 25S ribosomal subunit, respectively, as seen in Figure 6B.

The effect that short sRNA sequences are handled better by RNAplfold can also be seen in Figure 7. Here, we have categorised all nucleotides participating in RNA secondary structure or RNA-RNA interaction into three types: ‘only *cis*’, ‘only *trans*’, or ‘*cis & trans*’, as shown in Figure 7A. Figure 7B shows the histogram of accessibilities for all nucleotides per class, for the whole sRNA dataset (calculated with *W* = 240 and *L* = 160, as used in (31)). ‘Only *cis*’ nucleotides generally have very low accessibility, ‘only *trans*’ nucleotides generally have high accessibility, and ‘*cis & trans*’ also have generally low accessibilities (even though not as pronounced as ‘only *cis*’ nucleotides), all in agreement with the finding that estimated accessibility values agree well with the reference structure for sRNAs.

**Figure 7.**
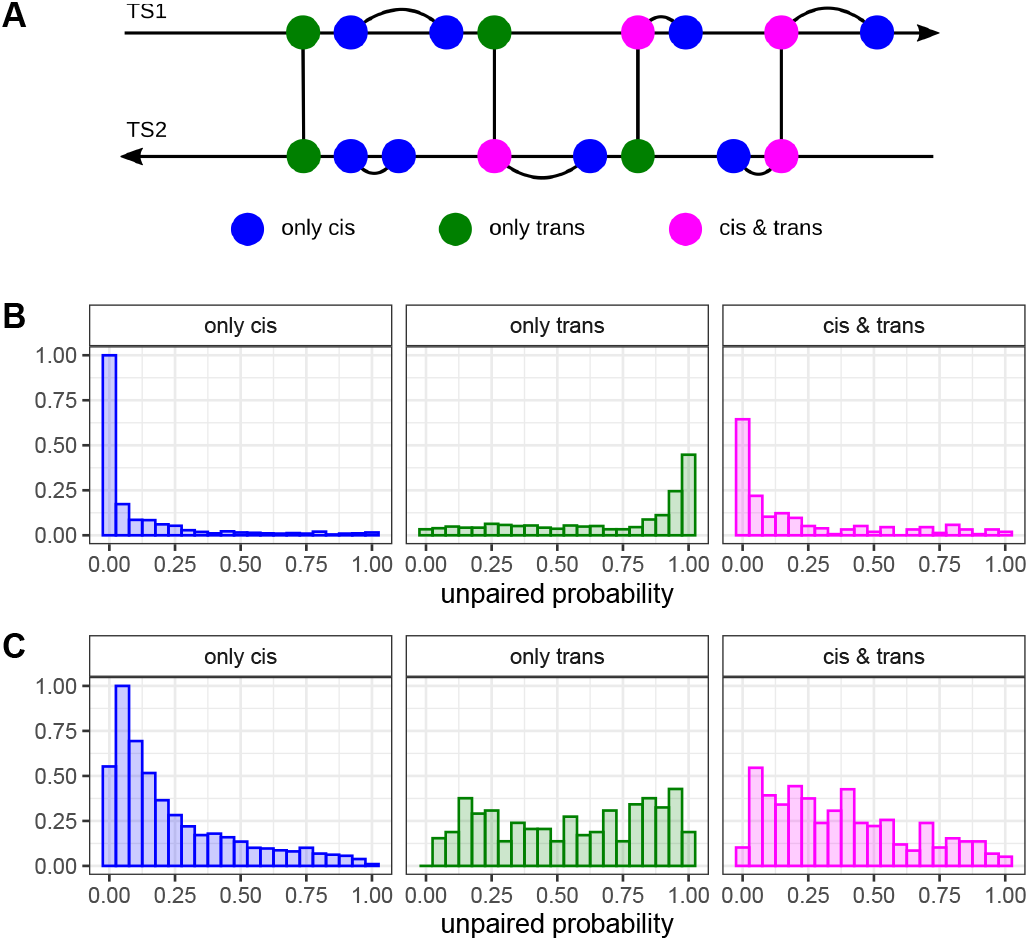
(A) Nucleotide classification, TS = transcript. (B) Unpaired probability (‘accessibility’) histograms for all sRNAs, for different classes according to (A). (C) Unpaired probability histograms for rRNAs, for different classes according to (A). Accessibility profiles have been generated with RNAplfold, settings *W* = 240, *L* = 160, as used in (31).

For the much longer transcripts of the ribosomal RNAs 18S and 25S sequences, on the other hand, Figure 7C shows that the accessibility distributions for the different types of nucleotides are much less pronounced: for ‘only *cis*’ nucleotides, accessibility is mainly low but there are also many nucleotides with medium or high accessibilities, for ‘only *trans*’ accessibilities are almost uniform and for ‘*cis & trans*’ there is only a very slight tendency towards lower accessibilities.

These findings are in line with early observations by Morgan & Higgs from 1996 (54) who found that the MFE RNA secondary structures predicted using the thermodynamic approach are typically in good agreement with the known, biologically functional RNA structures for transcripts shorter than around 200 nt. For longer transcripts, the agreement typically decreases with increasing transcript length. RNAplex authors discuss this also in their recent paper (55). Back then, Morgan & Higgs hypothesised that this disagreement is likely to be due to effects of co-transcriptional kinetic folding on RNA structure formation *in vivo*. We could show in 2013 that a minor modification of the typical thermodynamic approach to RNA structure prediction which takes the overall effect of co-transcriptional folding into account can significantly increase the prediction accuracy for long sequences (of more than 1000 nucleotides length) (56, 57, 58).

Is it thus not surprising that the estimation of accessibilities for sRNAs works fairly well (their sequence length is mostly below 200, with a maximum at 239), and that there is no improvement for larger window sizes in the rRNA accessibility prediction. RNA-RNA interaction prediction programs take this into account by choosing relatively small windows and maximum base pair distance as default values (IntaRNA: *W* = 150, *L* = 100, RNAplex: *W* = 240, *L* = 160). To be fair, one has to keep in mind that the accessibility profile is an ensemble property based on all possible pseudoknot free RNA structure conformations in all windows of size *W*, and is thus by construction not supposed to reflect (only) the MFE structure. Nevertheless, the maximum *Q* which can be reached for rRNAs (for any value of *W*) is fairly low and the unpaired probabilities frequently in disagreement to the known structure, as seen in Figure 5B.

We conclude that the estimation of accessibilities based on the thermodynamic model works only reasonably well for rather short molecules up to ≈200.

### The performance of RNA-RNA interaction prediction software is independent of the quality of the estimated accessibility profiles

We employed IntaRNA and RNAplex without and with accessibilities estimated by RNAplfold using different settings, to investigate if there is a correlation between the quality of the accessibility estimation *Q* and the prediction performance. As both RNA-RNA interaction prediction programs claim that the notion of accessibility is key to their superior predictive performance, we would expect better predictions to be due to more accurate estimation of the true underlying accessibilities. For this investigation, we used the accessibility settings given as defaults in the programs RNAplfold and IntaRNA, and those given for RNAplex in (31). Note that capitalised names (RNAplex, IntaRNA) refer to the prediction program which was used, while normal names (RNAplex etc.) refer to the default accessibility settings suggested to be used with the respective program. Additionally, we used two different settings ‘custom_1’ and ‘custom _2’ derived from the analysis of *Q* for all four classes of molecules. Settings which were used to generate sRNA/mRNA accessibilities are displayed in Table 1, and settings used to generate snoRNA/rRNA accessibilities are shown in Table 2.

**Table 1.** Accessibility settings for predictions for the sRNA-mRNA dataset. *S* is total sequence length. *q* is the query sequence (sRNA), *t* the target sequence (mRNA). *W* denotes the width of the sliding window in nucleotides, *L* is maximal base pair span.

| setting | $W_q$ | $L_q$ | $W_t$ | $L_t$ |
| --- | --- | --- | --- | --- |
| RNAP fold<sup>a</sup> | 70 | 70 | 70 | 70 |
| IntaRNA<sup>b</sup> | 150 | 100 | 150 | 100 |
| RNAp lex<sup>c</sup> | 240 | 160 | 240 | 160 |
| custom_1 | $S$ | $S$ | $0.25S$ | $0.25S$ |
| custom_2 | $S$ | $S$ | $0.75S$ | $0.75S$ |

<sup>a</sup>Default settings for RNAPL FOLD.
<sup>b</sup>Default settings for INTARNA ( $W = \text{acc}W$ , $L = \text{acc}L$ ).
<sup>c</sup>Settings from (31).

**Table 2.**
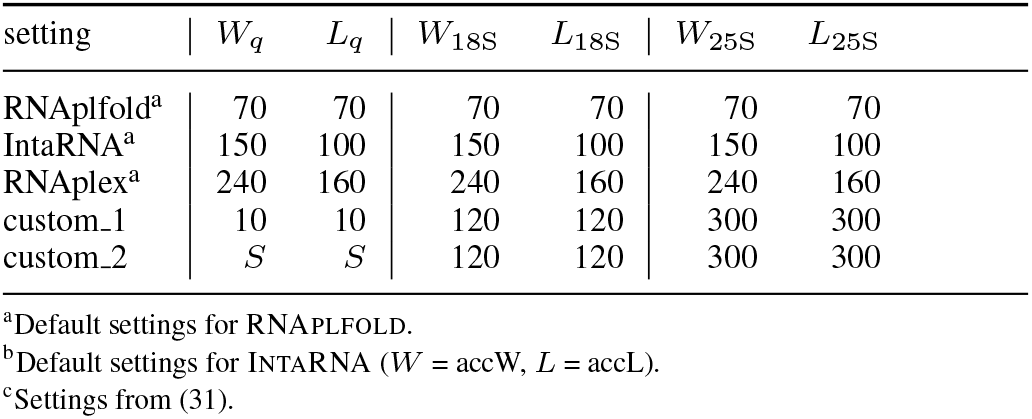
Accessibility settings for predictions for the snoRNA-rRNA dataset. *S* is total sequence length. *q* is the query sequence (snoRNA), targets are 18S and 25S ribosomal RNA. *W* is the sliding window size, *L* the maximal base pair span.

Figure 8 and Table 3 show the prediction performance of IntaRNA for the sRNA-mRNA dataset, as F1 score calculated based on the MFE prediction. What is noticeable is that without accessibilities, the performance is very bad (mean F1 0.08). Only in a few cases, the algorithm is able to identify several correct base pairs. Using accessibility with a small window length and maximum corresponding base pair span, *W* = 70 and *L* = 70 in the ‘RNAplfold’ setting, the mean F1 for the sRNA-mRNA dataset increases to 0.36. Increasing *W* and *L* further (‘IntaRNA’ setting) also increases the mean performance to 0.48. The three settings with the largest *W* and *L* (‘RNAplex’, ‘custom_1’, ‘custom_2’) yield very similar average performance values of 0.51 to 0.55. The prediction performance of RNAplex on the same dataset with the same accessibility settings is shown in Supplementary Material Figure S5. Mean F1 values are similar to IntaRNA, only for the settings ‘RNAplfold’ and ‘IntaRNA’, it is better by ≈ 0.05.

**Figure 8.**
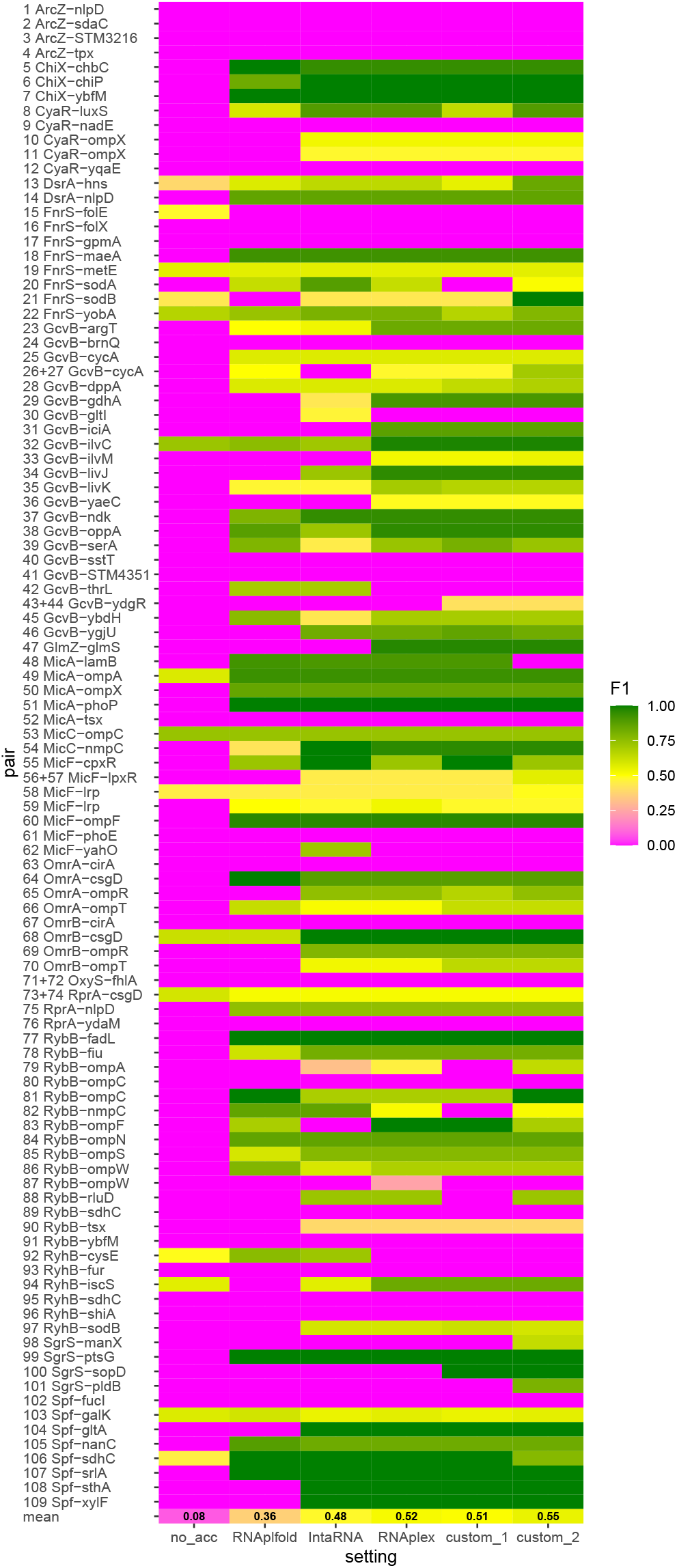
F1 performance score for the sRNA-mRNA dataset for predictions with IntaRNA for different accessibility settings. “no_acc” means no accessibility profiles where used. All other setting names correspond to Table 1. In the last row, the mean F1 value is given.

**Table 3.**
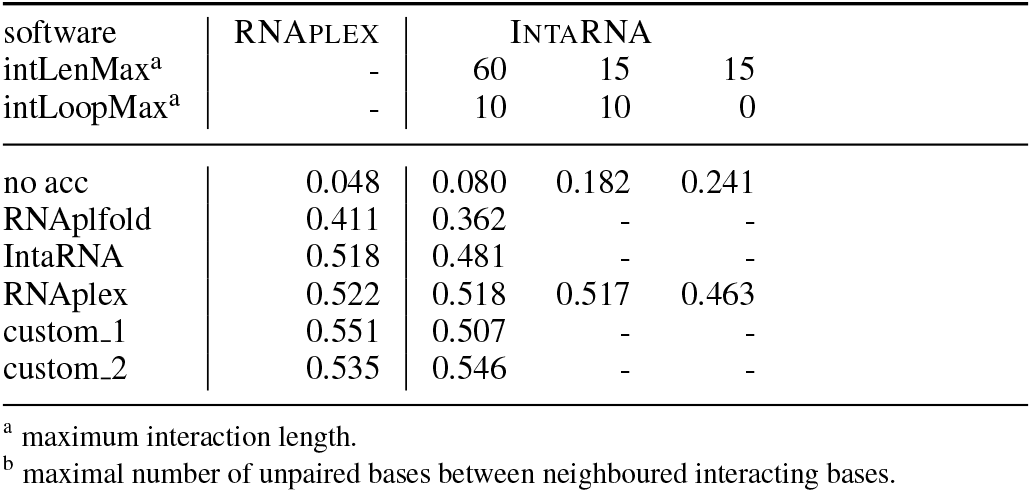
Predictions on sRNA-mRNA dataset, mean F1 values.

Table 5 shows the Pearson correlation coefficients *r* between the accessibility quality for the sRNAs *Q*_sRNA_ and the sensitivity, PPV and F1 score for the whole dataset, for each setting. All *r* values are positive, which we had expected, assuming that a better estimation of the underlying accessibility profile, i.e. a higher value of *Q*_sRNA_, yields better predictions, but have values generally below 0.2. For predictions by IntaRNA with settings with large *W* (RNAplex, custom_1, custom_2), *r* is only 0.07.

**Table 4.** Predictions on snoRNA-rRNA dataset, mean F1 values.

| software | RNAPLEX | INTARNA |  |  |
| --- | --- | --- | --- | --- |
| intLenMax<sup>a</sup> | - | 60 | 15 | 15 |
| intLoopMax<sup>b</sup> | - | 10 | 10 | 0 |
| no acc | 0.111 | 0.211 | 0.639 | 0.704 |
| RNAP fold | 0.494 | 0.522 | - | - |
| IntaRNA | 0.707 | 0.554 | - | - |
| RNAp lex | 0.691 | 0.594 | 0.761 | 0.775 |
| custom_1 | 0.390 | NA<sup>c</sup> | - | - |
| custom_2 | 0.707 | 0.581 | - | - |

<sup>a</sup> maximum interaction length.
<sup>b</sup> maximal number of unpaired bases between neighboured interacting bases.
<sup>c</sup> in INTARNA, $W$ has to be $\geq \text{intLenMax}$ (which was 60), so it was not possible to run it with these settings

**Table 5.** Predictions on sRNA-mRNA dataset, Pearson correlation coefficient *r* for *Q*_sRNA_ and Sens, PPV, F1, respectively.

| software → | RNAPLEX |  |  | INTARNA |  |  |
| --- | --- | --- | --- | --- | --- | --- |
| setting | $r_{\text{Sens}}^a$ | $r_{\text{PPV}}$ | $r_{\text{F1}}$ | $r_{\text{Sens}}$ | $r_{\text{PPV}}$ | $r_{\text{F1}}$ |
| RNAPfold | 0.12 | 0.12 | 0.12 | 0.12 | 0.12 | 0.12 |
| IntaRNA | 0.12 | 0.11 | 0.12 | 0.14 | 0.16 | 0.14 |
| RNAPlex | 0.12 | 0.10 | 0.11 | 0.11 | 0.07 | 0.07 |
| custom_1 | 0.21 | 0.16 | 0.19 | 0.12 | 0.04 | 0.07 |
| custom_2 | 0.14 | 0.15 | 0.15 | 0.09 | 0.06 | 0.07 |

<sup>a</sup> $r_{\text{Sens}} = r_{Q, \text{Sens}}$ etc.

Figure 9 and Table 4 show the prediction performance of IntaRNA for the snoRNA-rRNA dataset. Interestingly, predictions without accessibility have a relatively high mean F1 score of 0.21, opposed to 0.08 in the sRNA-mRNA dataset. Also for the other settings, the prediction performance is increased, the best setting being ‘RNAplex’ with 0.59. The prediction performance of RNAplex on the snoRNA-rRNA dataset with the same accessibility settings is shown in Supplementary Material Figure S6. With accessibility estimations, RNAplex performs systematically better than IntaRNA, reaching mean F1 scores of 0.69 to 0.71 for settings ‘IntaRNA’, ‘RNAplex’ and ‘custom_2’. It is worth noting that for setting ‘custom_1’, where *W* = *L* = 10 (derived from the best *Q* values for snoRNA in Supplementary Material Figure S4A), the prediction performance for RNAplex is low (0.39), showing that the assumption of mostly unstructured snoRNAs is detrimental for the correct prediction of snoRNA-rRNA interactions. For IntaRNA, this setting is prohibitive, because *W* is directly coupled to the maximum interaction length intLenMax, i.e. *W* ≥ intLenMax, with intLenMax = 60.

**Figure 9.**
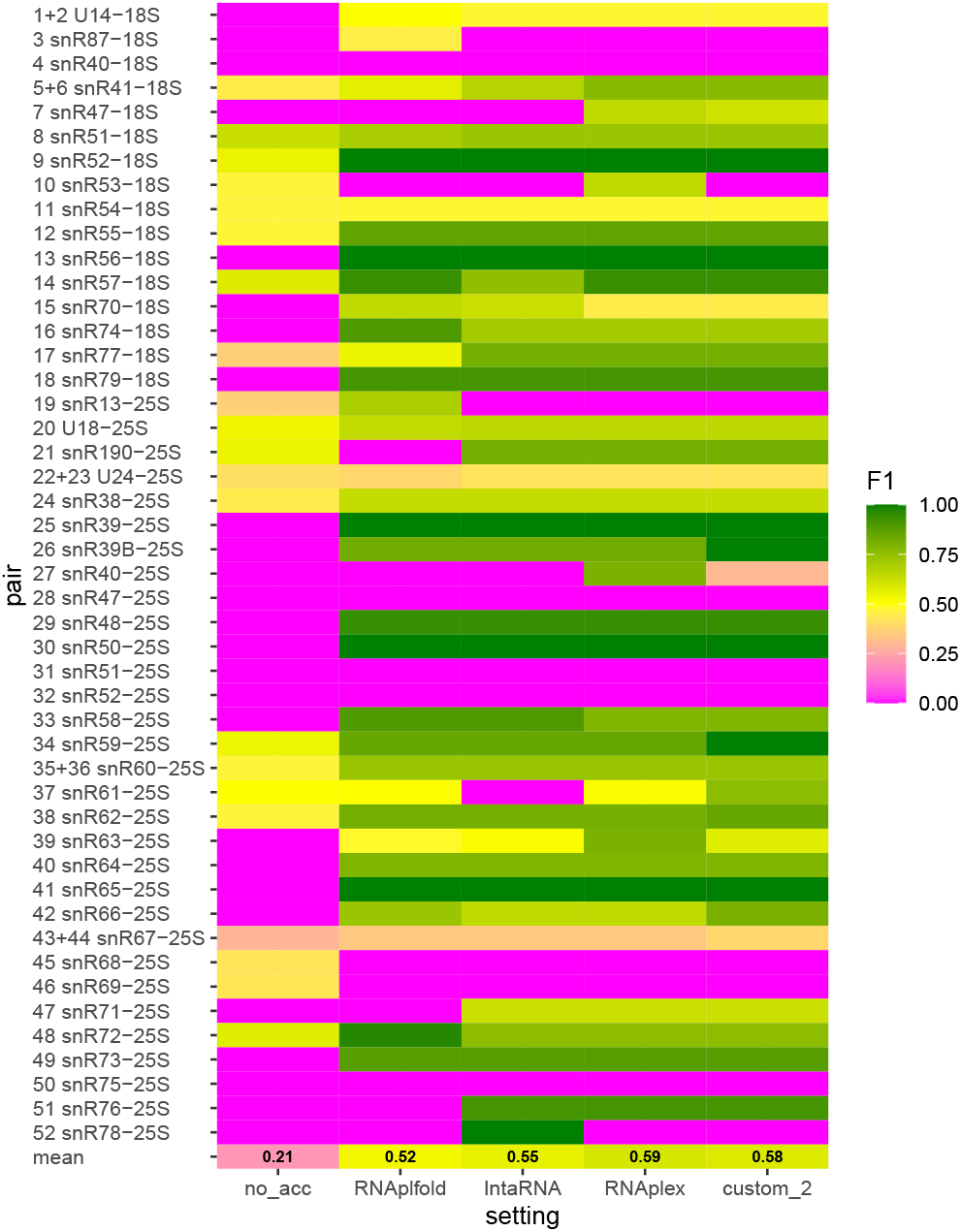
F1 performance score for the snoRNA-mRNA dataset for predictions with IntaRNA for different accessibility settings. “no_acc” means no accessibility profiles where used. All other setting names correspond to Table 2. In the last row, the mean F1 value is given.

Table 6 shows the pearson correlation coefficients *r* between *Q*_rRNA_ and sensitivity, PPV and F1 score for the snoRNA-rRNA dataset, for each setting. Again, the correlation *r* is very low in all cases (≤ 0.18), and even negative (≥ −0.18).

**Table 6.**
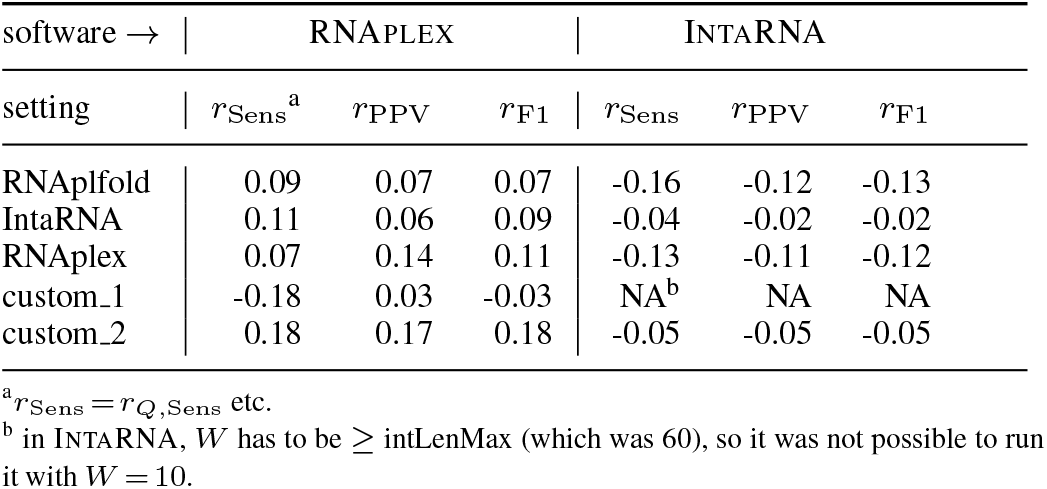
Predictions on snoRNA-rRNA dataset, Pearson correlation coefficient *r* for *Q*_rRNA_ and Sens, PPV, F1, respectively.

| software → | RNAPLEX |  |  | INTARNA |  |  |
| --- | --- | --- | --- | --- | --- | --- |
| setting | $r_{\text{Sens}}^a$ | $r_{\text{PPV}}$ | $r_{\text{F1}}$ | $r_{\text{Sens}}$ | $r_{\text{PPV}}$ | $r_{\text{F1}}$ |
| RNAPfold | 0.09 | 0.07 | 0.07 | -0.16 | -0.12 | -0.13 |
| IntaRNA | 0.11 | 0.06 | 0.09 | -0.04 | -0.02 | -0.02 |
| RNAPlex | 0.07 | 0.14 | 0.11 | -0.13 | -0.11 | -0.12 |
| custom_1 | -0.18 | 0.03 | -0.03 | NA<sup>b</sup> | NA | NA |
| custom_2 | 0.18 | 0.17 | 0.18 | -0.05 | -0.05 | -0.05 |

<sup>a</sup> $r_{\text{Sens}} = r_{Q, \text{Sens}}$ etc.
<sup>b</sup> in INTARNA, $W$ has to be $\geq \text{intLenMax}$ (which was 60), so it was not possible to run it with $W = 10$ .

To conclude, we find that there is only a negligible correlation between the quality of the accessibility estimation *Q* and the prediction performance of the two programs IntaRNA and RNAplex.

### One important overall effect of accessibility profiles is to prevent the thermodynamic model from forming too long interactions

Despite the shortcomings described above, the intriguing fact remains that, overall, RNA-RNA interaction prediction programs work much better when using accessibility profiles as opposed to not using them. To put it more drastically, see Figures 8 and 9, without accessibility profiles RNAplex and IntaRNA are almost completely incapable of identifying RNA-RNA interaction correctly. In order to understand the reasons for this, we looked into the details of the interactions predicted with and without accessibility estimation. What becomes apparent is that without accessibility estimation, both programs tend to predict very long duplexes, i.e. non-contiguous stretches of base pairs generally frequently interrupted by bulges or internal loops, which have very low energies according to the underlying thermodynamic models. It is important to note that these duplexes generally *do not contain the correct interaction site*. Figure 10C shows as a typical example the predicted interaction between snR56 and the 18S ribosomal subunit. It stretches over 57 nucleotides, close to the maximum interaction length set in IntaRNA, 60. When using accessibility profiles, see Figure 10A, an elongation of duplexes accross regions with low estimated accessibility is prevented by large energy penalties, leading to shorter duplexes altogether, and thus to better predictions, as seen in Figure 10B. In Figure 10D, we show that a similar effect can be achieved by simply reducing the default maximum interaction length (intLenMax) in IntaRNA to 15, without using accessibility profiles at all. Note that for RNAplex, it was not possible to set the maximum possible interaction length, so we have no data for RNAplex predictions in this case.

**Figure 10.**
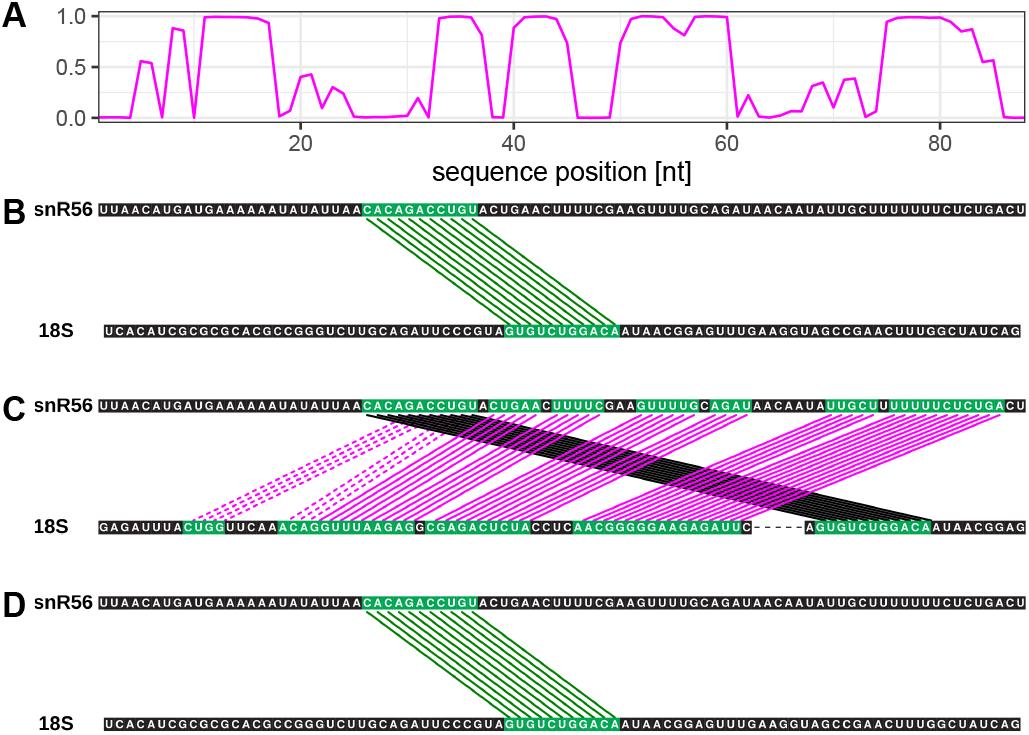
(A) (1-unpaired probability) for snR56. (B) Prediction with IntaRNA with accessibility setting ‘RNAplex’. (C) Prediction with IntaRNA without accessibility. (D) Prediction with IntaRNA without accessibility, but setting the maximal interaction length to 15. For (B) and (C), the maximum interaction length is 60. (B) to (D) color code: green: true positives, magenta: false positives, black: false negatives.

Figure 11 shows this effect for the complete snoRNA-rRNA dataset. Setting intLenMax to 15 recovers most of the correct predictions, without the need for utilising accessibility profiles. The mean F1 score increases significantly from 0.21 to 0.64, even better than with the best accessibility setting ‘RNAplex’ which lead to a mean F1 score of 0.59. Table 4 shows that by further decreasing the number of bulged/looped-out nucleotides (intLoopMax), this leads to an additional increase yielding 0.70. Combining accessibility profiles, intLenMax and intLoopMax leads to a further increase in performance, to 0.78.

**Figure 11.**
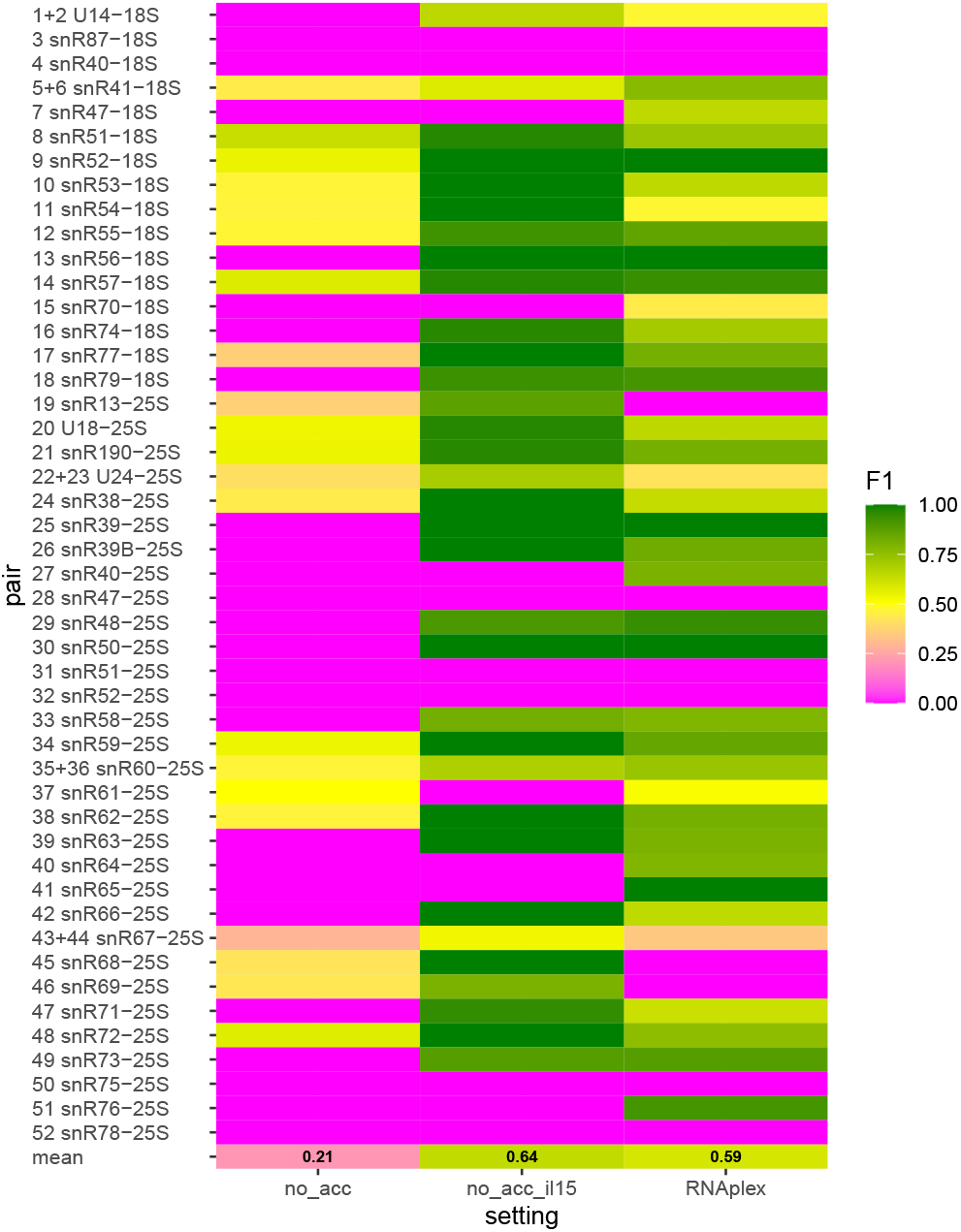
F1 performance score for the sRNA-mRNA dataset, predictions with IntaRNA without accessibility estimation (no_acc), without accessibility estimation, but with maximum interaction length set to 15 (no_acc_il15), and with accessibility estimation with setting ‘RNAplex’.

For the sRNA-mRNA dataset, which has different characteristics regarding interaction length and amount of bulged/looped-out nucleotides as can bee seen in Supplementary Material Figure S1, it is not as obvious how to boost the performance. Table 3 and Supplementary Material Figure S7 show that the performance without accessibility improves only slightly when setting intLenMax to 15, and intLoopMax to 0. When using these settings together with accessibility profiles, the prediction performance gets even worse, because many of the known interactions are longer than 15 and can no longer be found.

Overall, we thus conclude that one important overall effect of estimating accessibility profiles is to limit the length of duplexes. In biological classes with predominantly short, highly complementary interactions like the snoRNA-rRNA interactions, this effect is sufficient to boost the prediction performance, and can be mimicked by simply limiting the maximum permitted interaction length. In biological classes with more diverse interactions in terms of interaction length, and number of bulges/internal loops like sRNA-mRNA interactions, this effect is present, but not as influential for the performance. Here, a precise estimation of sRNA accessibility is essential for a good prediction performance.

## DISCUSSION

Predicting novel, functionally relevant *trans* RNA-RNA interactions *de novo*, i.e. based on sequence information only, remains one of the most intriguing open challenges in computational biology today. The best state-of-the-art methods today, RNAplex and IntaRNA, rely on the concept of accessibility to reach their superior predictive performance. This concept is based on the assumptions that (1) RNA-RNA interactions binding sites tend to be devoid of RNA secondary structure features, (2) before *trans* binding occurs, the two interacting transcripts are each in thermodynamic equilibrium and also not interacting with any other molecules inside the cell, (3) RNA-RNA interaction formation also follows the MFE principle, i.e. conflicting RNA secondary structure base pairs first have to be opened, penalising the RNA-RNA interaction energy, and (4) the two interacting transcripts are already fully transcribed at the moment of interaction.

As recent progress in the field of *trans* RNA-RNA interaction prediction has been scarce since the publication of the two state-of-the-art programs RNAplex in 2011 and IntaRNA 2008 (version 2 in 2017) and as there is still ample room for improving the predictive performance in the field, we have, for the first time, investigated the concept of accessibility and its underlying assumptions.

We find that known *trans* RNA-RNA interactions often overlap RNA structure features, contradicting assumption (1). This is the case for short sRNAs, even more so for long ribosomal subunits, and presumably also for mRNA structures. This is not overly surprising, given that RNA structure features and *trans* RNA-RNA interaction may affect the RNA transcript at different times of its cellular life. This is especially evident for snoRNA-rRNA interactions, which happen during ribosome biogenesis, before the mature ribosomal RNA structure is formed. All *cis* and *trans* basepairs required throughout each transcript’s life *in vivo* are encoded in the respective transcript’s sequence. The two state-of-the-art programs IntaRNA and RNAplex, however, have no way of knowing which of these *cis* and *trans* features are required *at the same point of time in vivo*. Our results clearly show that the notion of accessibility for improving the prediction of *trans* interactions has to be conceptually questioned as known *trans* interacting regions quite often overlap regions that are also known to interact in *cis* at some point of the transcript’s life *in vivo*.

The amount of conflicts between these *cis* and *trans* base pairs, however, does not have an impact on the prediction accuracy of the *trans* RNA-RNA interactions. This can have two reasons: either, accessibility profiles somehow intrinsically reflect possible *cis*-*trans* conflicts, or, for the region of interest, the exact accessibility profile is not the dominant factor to rank the predicted duplexes by their free energy. For sRNAs, the accessibility of nucleotides with *cis-trans* conflicts is generally low, similar to ‘only *cis*’ nucleotides. Also for long rRNAs, the conflicting nucleotides tend to have lower accessibilities, even though much less pronounced than ‘only *cis*’ nucleotides. In other words, in most cases, the accessibility-based approach disfavours conflicting nucleotides from being *trans* base-paired. This is in agreement with the way accessibility is being calculated, which is solely based on MFE estimations of RNA structure features of an isolated RNA molecule in thermodynamic equilibrium.

We defined a new measure *Q* to quantify the agreement between the estimated accessibilities and the known reference RNA secondary structure. RNAplfold does a fairly good job estimating the accessibility of short sRNAs, if optimal parameters are used. For long sequences like the 18S and 25S ribosomal subunits, however, the agreement between the true structure and the predicted accessibility profile is mediocre, showing that for long sequences, the MFE-based strategy is sub-optimal, even when limiting the maximum base pair span. This discrepancy may be due to *in vivo* effects such as co-transcriptional folding or the participation of other potential *trans* interaction partners such as proteins, both of which are being ignored by assumptions (2) and (4). Most strikingly, we find that performance with which *trans* RNA-RNA interactions are predicted does not correlate with the quality of correctly estimating the respective accessibilities.

Despite these results, both state-of-the-art programs for predicting *trans* RNA-RNA interactions perform much better when using estimated accessibilities, compared to not using accessibilities. This can be explained by the one important overall effect that the accessibility profiles have, namely the prevention of unreasonably long duplexes. The MFE strategy heavily encourages the prediction of long RNA-RNA interaction duplexes (until they reach close to the maximum permitted interaction length). Low accessibility and therefore high energy penalties at key positions make long duplexes energetically unfavourable compared to shorter duplexes, thereby increasing the prediction accuracy. We show that for the snoRNA-rRNA dataset, reducing the maximum interaction length (and the number of mismatches/bulges) without using accessibility estimates leads to a better performance than when including accessibility estimates. A combination of all three leads to the best performance for this dataset. For the sRNA-mRNA dataset, which has more heterogeneous characteristics in terms of interaction length and mismatches/bulges, this effect can also be seen, yet to a lower degree. Here, well estimated accessibility profiles are clearly beneficial.

We conclude that when predicting novel *trans* RNA-RNA interactions in a class where no validated interactions exist, the safest strategy is using accessibility profiles with the parameters we found to work best. If, however, there is additional knowledge about the characteristics of the *trans* RNA-RNA interaction in a biological class, such as the typical length or number of mismatches/bulges, this information can be leveraged to prevent predictions not fitting the criteria.

Without accessibility (and with default maximal interaction length), the MFE approach to RNA-RNA interaction prediction employed by IntaRNA and RNAplex simply does not work, because by construction the method tries to increase the duplex length as much as possible. With estimated accessibility profiles, even if they would perfectly reflect the true RNA secondary structure of the two transcripts, even this RNA-RNA interaction prediction approach would be limited, because we have shown that conflicting *cis-trans* base pairs frequently co-exist. In order to improve the field of *trans* RNA-RNA interaction prediction further, we therefore suggest to focus future research on devising comparative methods for the prediction of novel *trans* RNA-RNA interaction.

The existing comparative methods (27, 28), however, require a fixed input alignment for each of the two transcripts of interest and are known to have a prediction performance which strongly depends on the alignment quality. As additional challenge, *trans* base pairs typically do not exhibit the same level of covariation as RNA structure features and are thus conceptually harder to detect using the established computational strategies for predicting RNA secondary structures in a comparative way. In order to significantly improve upon the current state-of-the-art in predicting *trans* interactions, we will thus not only require comparative methods that operate in an alignment-free manner, but also conceptually novel strategies to distinguish between evolutionarily conserved *cis* and *trans* base pairs. Even then, one remaining conceptual challenge needs to acknowledge the humbling fact that different biological classes of transcripts may require different functional RNA structures and *trans* RNA-RNA interactions throughout their cellular life *in vivo*. We can reasonably expect that some information on these functionally relevant *cis* and *trans* features is encoded in each transcript in question. Yet, which of these features is expressed when *in vivo*, is also determined by the particular details of the complex *in vivo* environment at a given point of time and space. Right now, none of the computational methods conceptually aim to disentangle the conserved *cis* and *trans* features into self-consistent configurations of mutually compatible *cis* and *trans* features that could be expressed at different points in time. As we discover more complexities *in vivo*, we may thus be required to expand the notion of alternative RNA structure expression (59) to the notion of alternative *trans* RNA-RNA interactions.

